# Contraction type dependence of serial sarcomere addition following resistance training

**DOI:** 10.64898/2026.09.29.755478

**Authors:** Amelia Rilling, Alexander M Zero, Alexandra Q Kirkup, Maxum Stratford, Kathryn Chapman, Avery Hinks, Geoffrey A. Power

## Abstract

Longitudinal muscle growth occurs through the addition of sarcomeres in series, a process termed serial sarcomerogenesis. To investigate contraction mode dependence of sarcomerogenesis in Sprague-Dawley rats (n=36, 14-18 weeks, ∼468g), we compared the effects of isometric, eccentric, and concentric training (4-6 weeks; 3×week) on morphological and mechanical adaptations of the plantar flexors. Isometric training was performed at a joint angle corresponding to the plateau nearing the descending limb of the force–length relationship, eccentric contractions began at a shorter muscle length and lengthened to this same joint position, and concentric contractions started at this lengthened position but then actively shortened. Owing to the rapid regulation of serial sarcomere number the 4-and 6-week timepoints were collapsed. While force output increased across all training modes (+16%; p<0.05), longitudinal muscle growth was regulated by contraction type. Both isometric and eccentric training were highly effective at increasing serial sarcomere number when optimal loading conditions (high forces at longer lengths) are provided (+7-9%; p<0.05). Concentric training, even when initiated at long lengths with high forces was not an effective stimulus for longitudinal remodelling (+2%; p>0.05). Therefore, these findings identify the mechanical loading environment associated with isometric and eccentric training as a key driver of serial sarcomerogenesis.

## Introduction

Skeletal muscle has the remarkable ability to undergo architectural remodelling in response to changes in the environmental loading condition imposed upon it. Remodelling occurs in both radial (muscle thickness) and longitudinal (along the length of the fibre) directions. Radial growth is driven primarily by the addition of myofibrils in parallel to one another, which increases fibre cross-sectional area. Conversely, longitudinal muscle growth occurs through the addition of sarcomeres in series along a myofibril, thereby increasing muscle fibre length, a process termed “serial sarcomerogenesis”^1^. Radial growth and longitudinal growth appear to be regulated by different mechanisms^2^, indicating that the type of training modality can impact the location and extent of sarcomere addition. For longitudinal growth there is little debate regarding the large magnitude increases in serial sarcomere number (SSN) following casting interventions at long muscle lengths^3^, however contraction mode dependent increases in SSN following training is widely debated and less clear^4–6^.

To date, only one study has examined SSN adaptations following isometric training. Uçar *et al.* (2024) reported a 12% and 32% increase in SSN after eight weeks of isometric training at intermediate and long muscle-tendon unit (MTU) lengths, respectively – highlighting the length-dependence of SSN adaptation^7^. This magnitude of SSN increase is higher than reports following eccentric training. SSN adaptations following eccentric training, whether that be downhill running (eccentric biased)^8–10^ or controlled dynamometry training^11–14^ can range from 0 to +11%. Eccentric overload training (maximal stimulus throughout the eccentric phase) has shown up to +17% increases in SSN^15^. Though not without criticism^4^, a recent paper^5^ suggested eccentric training was not an effective stimulus for serial sacromerogenesis. However, the data included into the systematic and meta-analytic review has been criticized^4,5^ and overall, more experimental data is warranted to help draw informed conclusions.

Conversely, the use of uphill treadmill training in rodents (biasing concentric contractions) yields inconsistent findings. Following training, some studies report a reduction in SSN (−4 to −11%)^13^, while others show no significant changes (+3 to −5%; ^8,16,17^). These discrepancies across concentric training (uphill walking/running) studies may reflect the variability of treadmill exercise. Overall, eccentric and isometric training appear to elicit increases in serial sarcomere number (SSN), whereas findings following concentric training are less consistent. These observations suggest that training-induced sarcomerogenesis may be dependent on contraction mode. However, this possibility has not been systematically investigated through direct comparisons of different contraction modes using a standardized experimental approach within the same laboratory. Such an approach is necessary to minimize methodological variability between studies and to determine whether differences in sarcomerogenesis can be attributed to contraction mode rather than to differences in experimental design, training protocols, or measurement techniques. Dynamometer-based resistance training offers a standardized approach to investigate contraction-type dependent adaptations by controlling joint range of motion, excursion, and activation timing, thereby producing a more optimal and consistent loading environment.

Eccentric contractions and isometric contractions at longer muscle lengths may preferentially stimulate serial sacromerogenesis by producing high forces at long muscle lengths, increasing mechanical loading of key mechanosensors such as titin and integrin-FAK complexes^1^, demonstrating that the type of contraction mode and stimulus may influence the type of remodelling. As serial sarcomere number influences the force–length relationship of skeletal muscle, understanding whether sarcomerogenesis is dependent on contraction mode is critical for determining how training programs can be designed to optimally modulate muscle architecture and functional capacity.

Here we investigated SSN and mechanical function of the rat plantar flexors following upwards of 4 weeks of resistance training. Either eccentric (active muscle lengthening), concentric (active muscle shortening), or isometric (active fixed-end contraction) training was performed on a dynamometer controlling for joint excursion, activation timing, and velocity, thus allowing for comparisons across training modalities. Multiple testable hypotheses emerged from a recent systematic review and meta-analysis of sarcomerogenesis in animal models^5^: (**i**) Muscle excursion has little to no effect, but end-contraction fibre length and force is the driving stimulus promoting the largest increase in SSN - therefore in the present study we should expect the largest magnitude of SSN increases following maximal isometric training as compared with sub-maximal eccentric training ending at the same muscle length. (**ii**) the authors^5^ suggest that eccentric exercise is not an effective promotor of serial sarcomerogenesis in animal models and *therefore* we should not expect a meaningful increase in SSN following eccentric training. Finally (**iii**), there are no available SSN data on concentric resistance training, and while there is generally a loss of SSN with uphill running, SSN should increase following concentric training if concentric contractions are initiated with high forces at long muscle lengths^5^. A secondary aim was to assess the concomitant adaptations in mechanical function following training.

## Results

### Data Exclusion

One rat was excluded from the concentric group mechanical data due to health complications during Post-training testing. The concentric sample size was adjusted to n=11 for mechanical data and n=12 for morphological data. The total sample size for mechanical data was n=35 and for morphological data was n=36.

### Isometric Training Mechanical Results

pre- to post-training, torque increased ∼18% at 90° and ∼23% at 70° with an effect of training (*p*<0.001; *η*^2^=0.074; Figure 3) and angle (*p*=0.002: *η*^2^=0.358; Figure 3) due to greater torque at 70°, but no training × angle interaction (*p*=0.182; *η*^2^=0.08). Passive tension had an effect of training (*p*<0.001; *η*^2^=0.696) and an interaction of training × angle (*p*=0.016; *η*^2^=0.238; Figure 3), however post hoc testing revealed both angles demonstrated comparable relative improvements following training (77% vs 74%) suggesting the interaction may reflect small differences in the magnitude of change rather than a substantially different training response. Additionally, there was an effect of angle (*p*<0.001; *η*^2^=0.686) as 70° generated greater passive tension. Peak power increased ∼23% with training (*p*=0.014; *η*^2^=0.437; Figure 3) driven by an ∼38% increase in torque at peak power (*p*<0.001; *η*^2^=0.904) despite ∼11% reductions in angular velocity at peak power (*p*=0.041; *η*^2^=0.328; Supplemental Figure 4), indicating a shift in peak power to higher loads following training.

**Figure 1.**
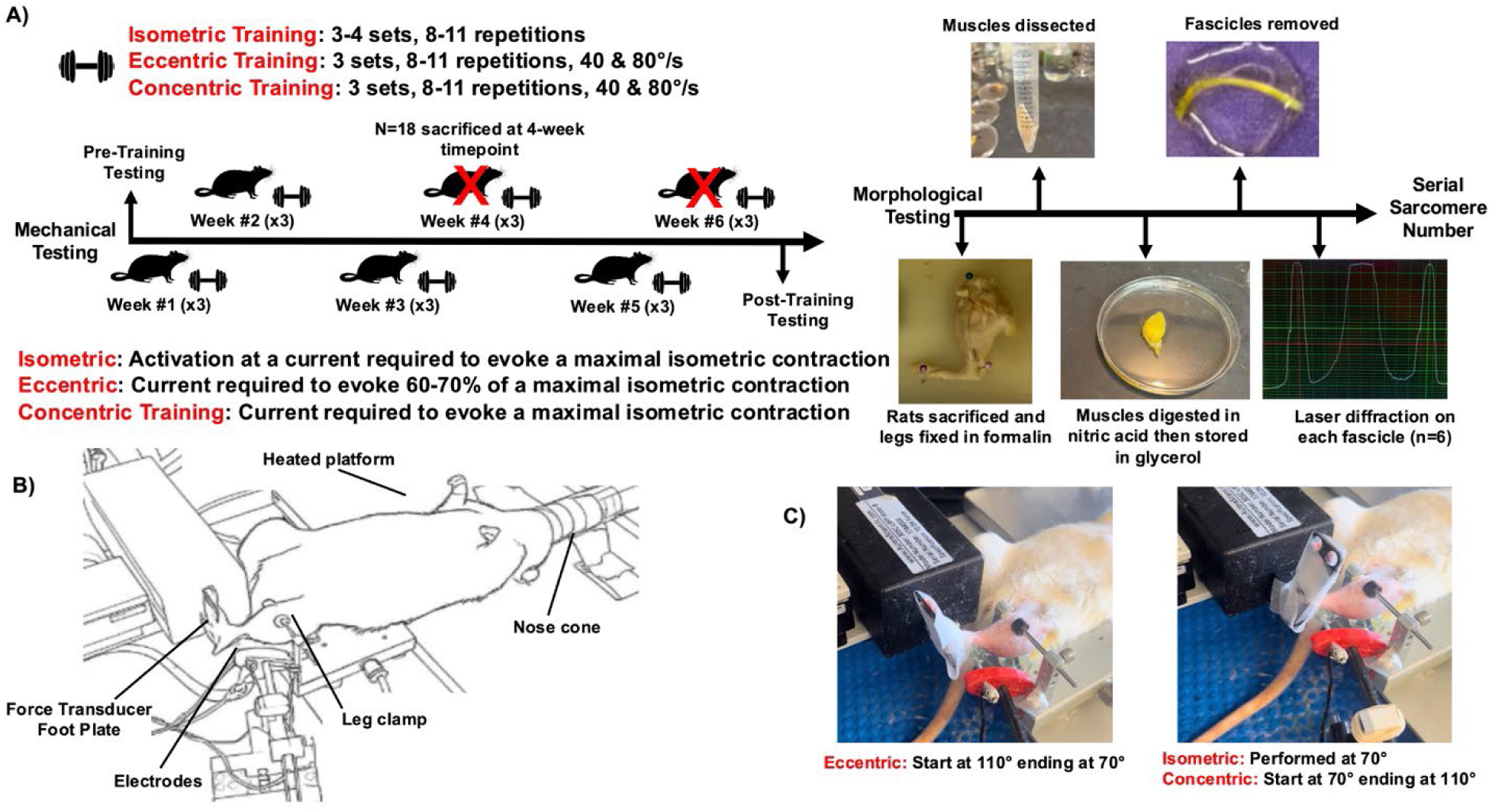
Experimental timeline demonstrating mechanical testing, training and morphological data collection. Male rats (n=36) underwent 4 to 6 weeks of isometric, eccentric or concentric training. Mechanical testing was done pre and post 4 weeks or 6 weeks of training on the same leg. Rats were anaesthetized and placed supine on a heated platform (37°C) with a nose cone providing isoflurane. The left leg was secured with a leg clamp around the knee (150°), with the left foot secured with tape to a foot plate attached to a torque motor/transducer. Percutaneous stimulation electrodes were positioned over the plantar flexors as well as the tibial nerve to evoke contractions. The ankle is pictured at 90° with a range of motion of 70° to 110°. Following training and mechanical testing rats were sacrificed for morphological data collection to calculate serial sarcomere number, with the non-trained leg acting as the internal control.

**Figure 2.**
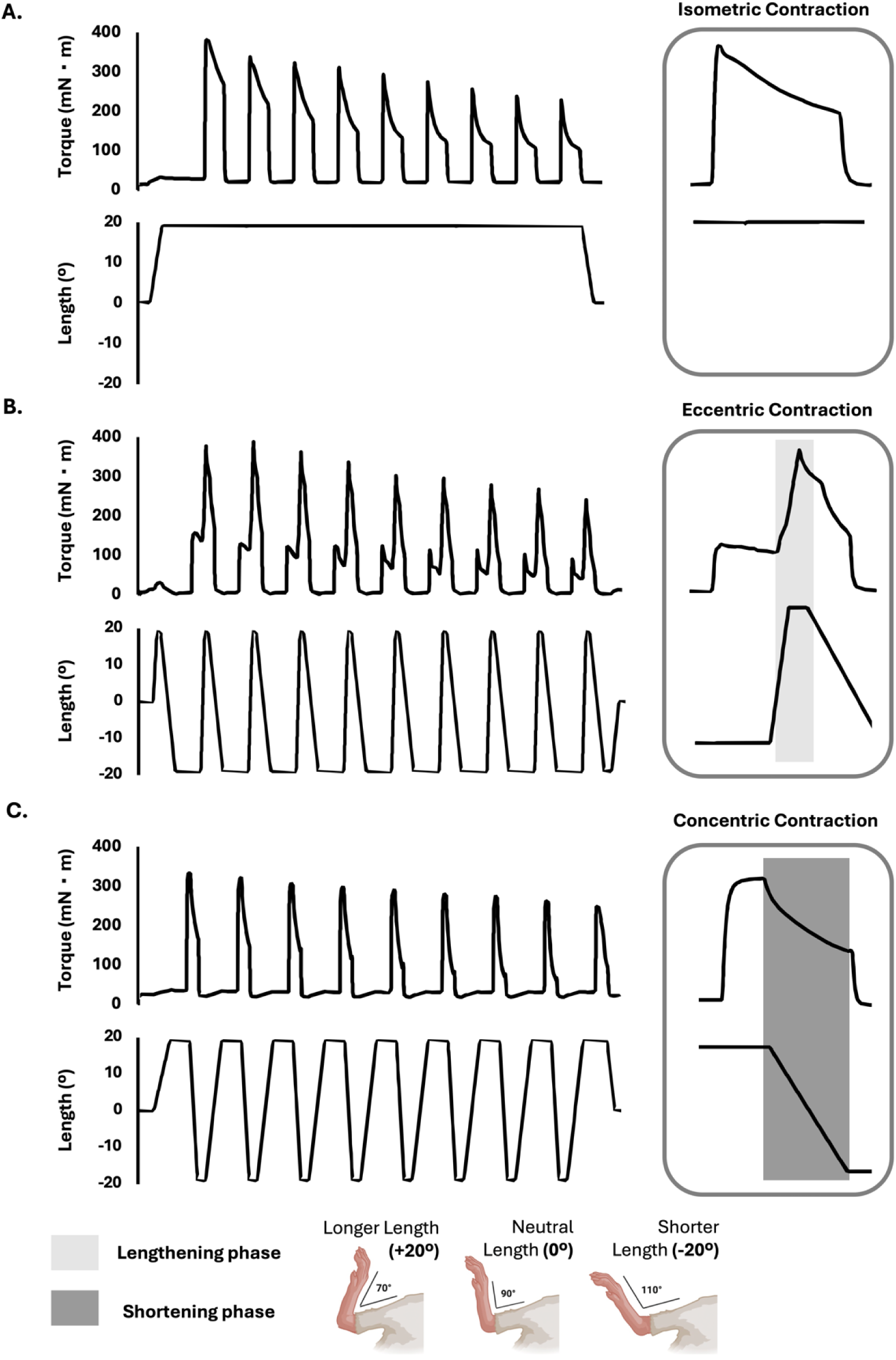
**Torque and position traces** of a training set for **A) isometric, B) eccentric and C) concentric training.** Isometric training was performed at 70°, training range of motion for concentric was 70° to 110° and reversed for eccentric. Activation level was maximal for isometric and concentric, and was set to 60-70% maximal tetanic isometric torque for eccentric.

**Figure 3.**
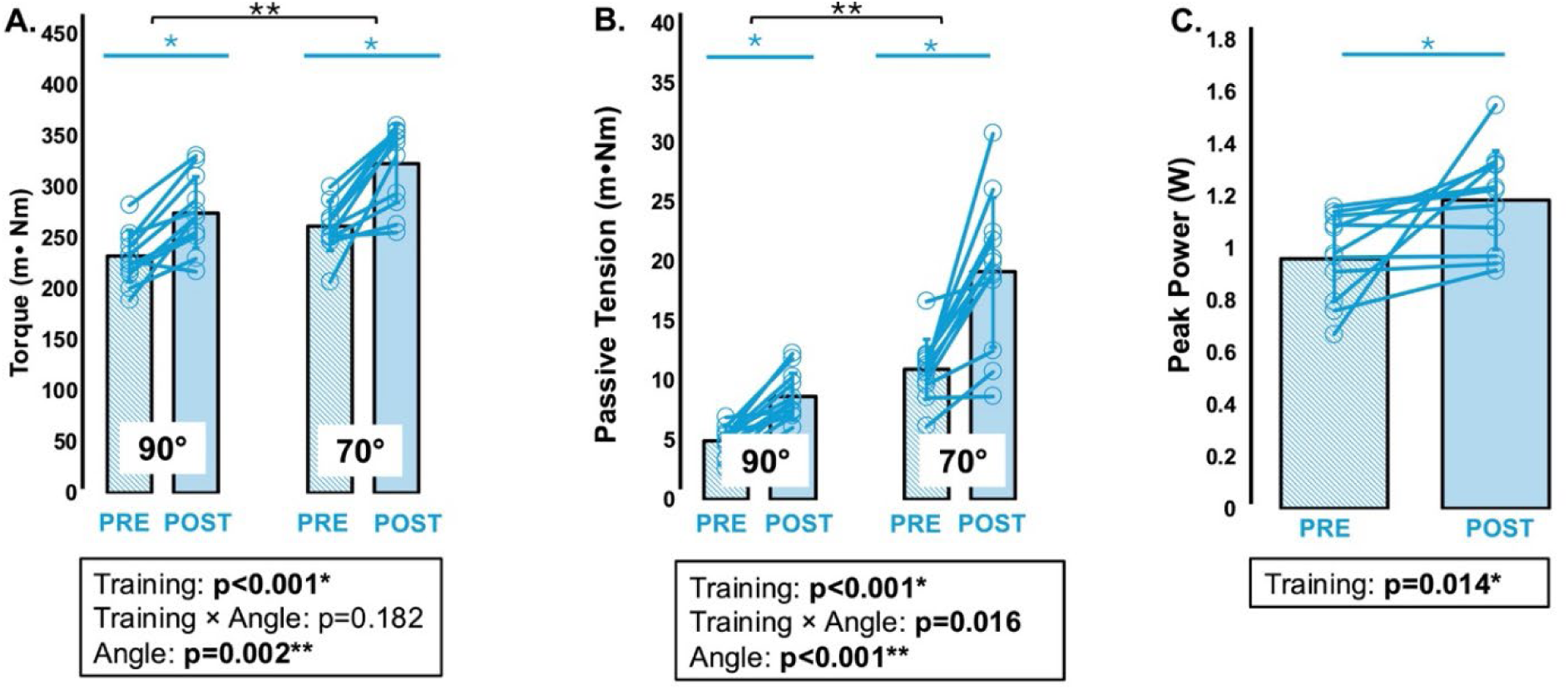
Isometric Training. Differences in plantar flexor **A) Torque at 90° (shorter muscle length) and 70° (longer muscle length)**, B) **Passive tension at 90° and 70° and C) Peak power** with isometric trained rats (n=12). Pretraining values are shown as blue lines and post training values are shaded blue. Statistical comparisons were performed using a two-way repeated measures ANOVA (Training ×Angle). Data are displayed as mean ± standard deviation. * Denotes significant differences.

### Isometric Training Morphological Results

Following isometric training, for the soleus there was a significant 9% increase in SSN (*p*<0.001, *η*^2^=0.720; Figure 4) leading to increases in fascicle lengths (Supplemental Figure 1) with no change in SL (Supplemental Figure 1). There was no change in soleus PCSA (*p*=0.893; *η*^2^=0.002; Figure 4), however soleus wet weight was trending towards an increase (*p*=0.064; *η*^2^=0.278; Figure 4) with training. There was no change in SSN in the MG (*p*=0.502; *η*^2^=0.042; Figure 4) nor FL or SL (Supplemental Figure 2). There was an ∼8% increase in wet weight with training (*p*=0.007; *η*^2^=0.503; Figure 4), and MG PCSA did not increase significantly with training (*p*=0.436; *η*^2^=0.056; Figure 4). The increased wet weight with no change in PCSA was likely owing to variability in FL changes and/or non-contractile tissue accumulation following training.

**Figure 4.**
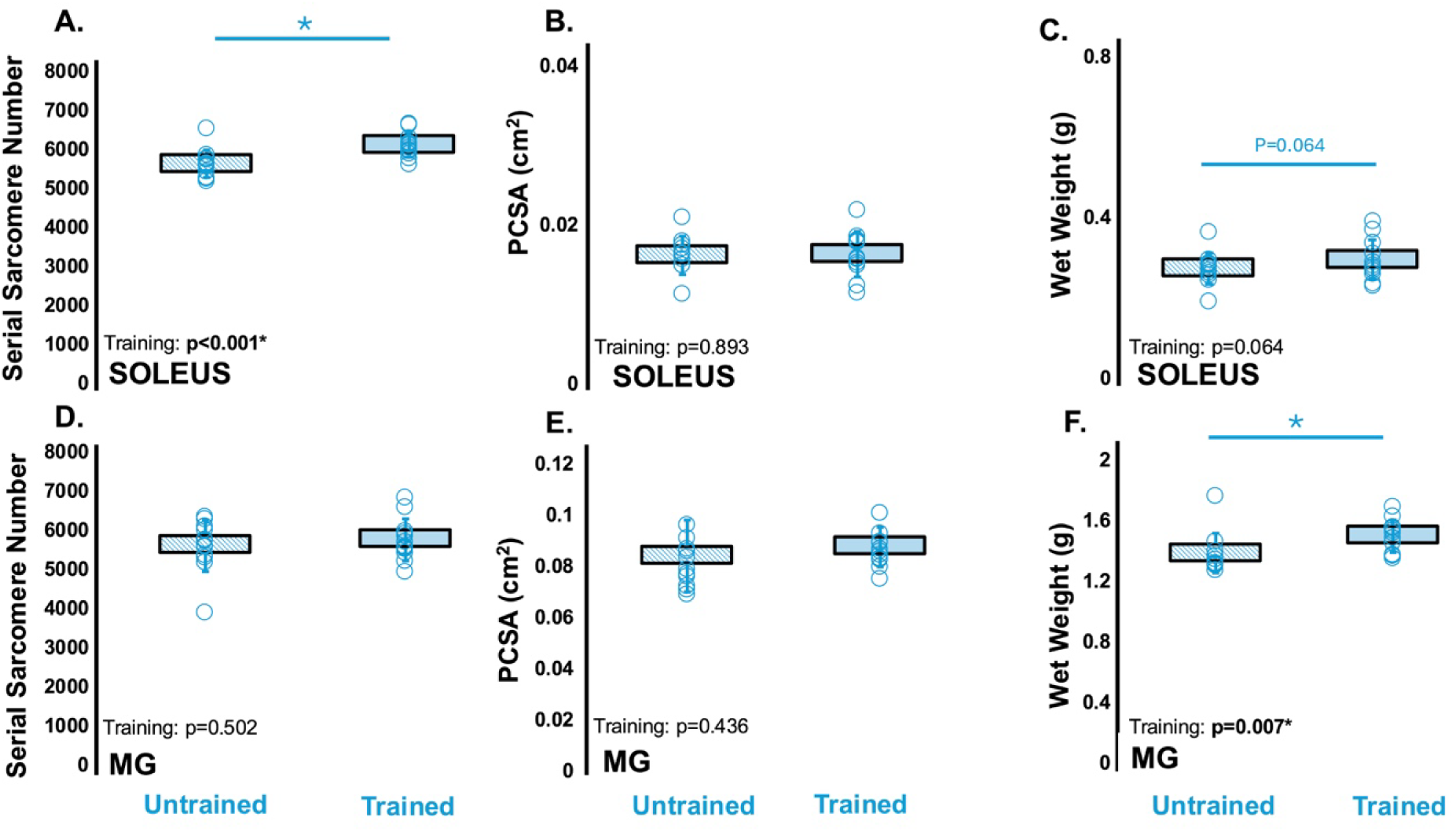
Isometric Training. **A) soleus SSN, B) soleus PCSA, C) soleus wet weight, D) MG SSN, E) MG PCSA and F) MG wet weight** following isometric training in male rats (n=12). The untrained (right leg) values are shown as blue lines and trained (left leg) values are shaded blue. Statistical comparisons were performed using a one-way repeated measures ANOVA (Training). Data are displayed as mean ± standard deviation. * Denotes significant differences.

### Eccentric Training Mechanical Results

There was an effect of training where torque at 90**°** increased ∼13% and at 70° increased ∼12% (p<0.001; η^2^=0.473; Figure 5). There was an effect of angle (p<0.001; η^2^=0.521) as there was greater torque at 70**°**. Passive tension had an ∼1% increase at 90**°** and an ∼38% increase at 70**°** (p=0.012; η^2^=0.255; Figure 5), with an interaction of training × angle (p=0.014; η^2^=0.245) due to greater increases in passive tension with training at 70**°**. There was an effect of angle (p<0.001; η^2^=0.756) driven by greater torque at 70**°**. However, peak power did not increase following training (p=0.926; η^2^=0.001; Figure 5) despite an increase in torque at peak power (p=0.028; η^2^=0.368) and no change in angular velocity at peak power (p=0.082; η^2^=0.250; Supplemental Figure 4).

**Figure 5.**
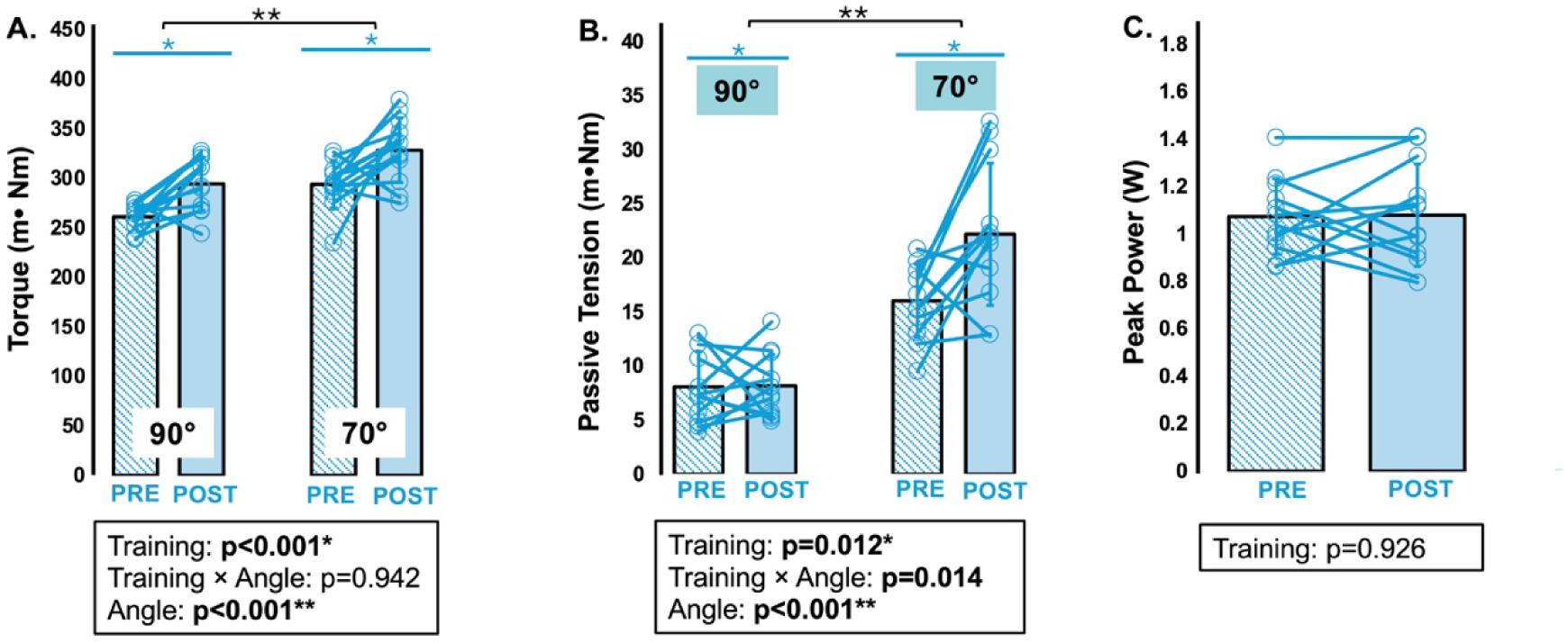
Eccentric Training. Plantar flexor **A) Torque at 90° (shorter muscle length) and 70° (longer muscle length), B) Passive tension at 90° and 70° and C) Peak power** following eccentric training in male rats (n=12). Pretraining values are shown as blue lines and post training values are shaded blue. Statistical comparisons were performed using a two-way repeated measures ANOVA (Training ×Angle). Data are displayed mean ± standard deviation. * Denotes significant differences.

### Eccentric Training Morphological Results

Following eccentric training there was a significant 7% increase in soleus SSN (p<0.001; η^2^=0.732; Figure 6) leading to increases in FL (Supplemental Figure 2) and no change in SL (Supplemental Figure 2). Soleus PCSA did not change with training (p=0.331; η^2^=0.086; Figure 6). Additionally, there was no change in soleus wet weight (p=0.694; η^2^=0.015; Figure 6). SSN in the MG increased ∼7% with training (p=0.028; η^2^=0.366; Figure 6) along with an increase in FL (Supplemental Figure 2) and no change in SL (Supplemental Figure 2). MG PCSA and wet weight did not change with training (p=0.225; η^2^=0.131; Figure 6, p=0.443; η^2^=0.054; Figure 6).

**Figure 6.**
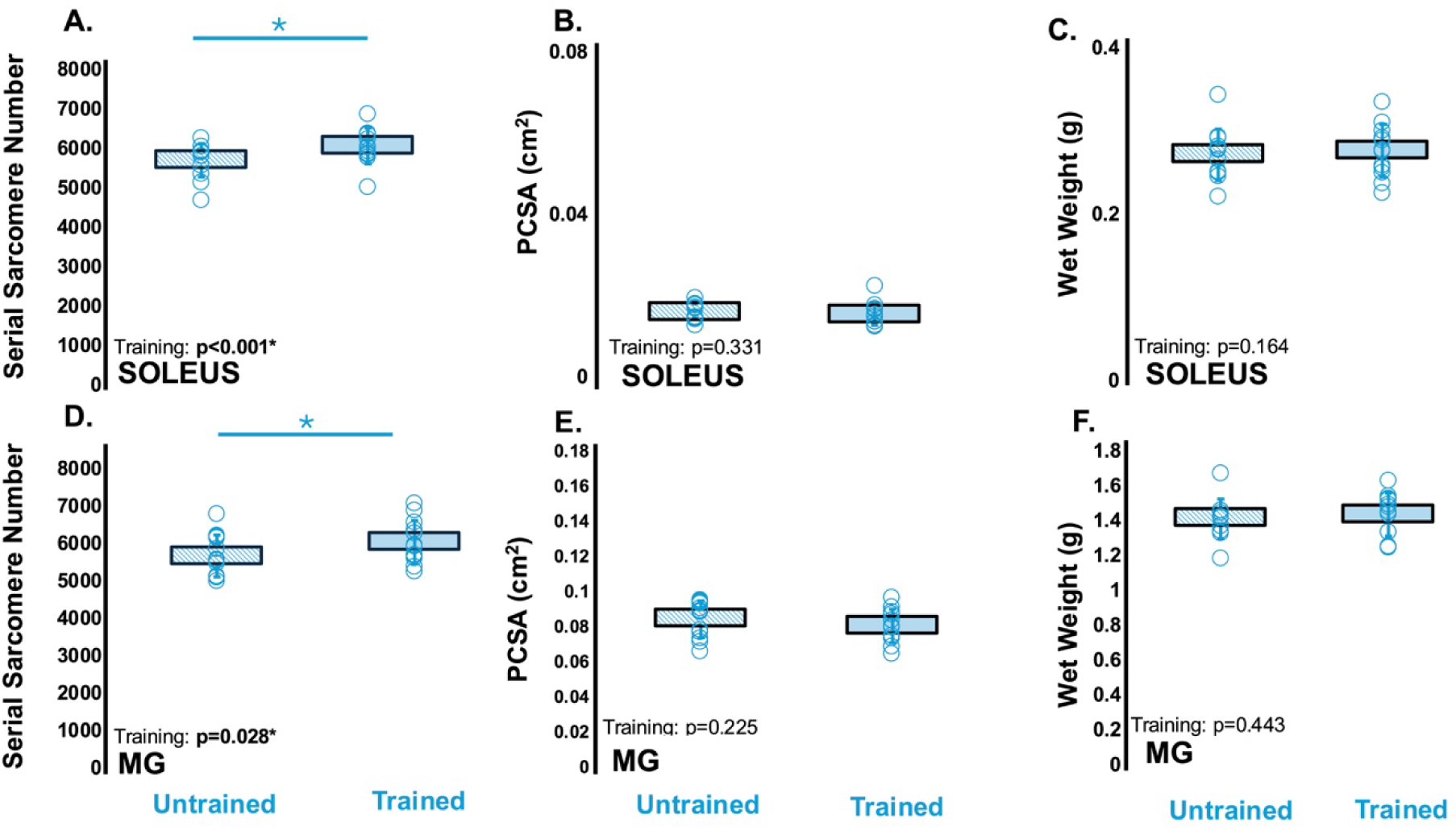
Eccentric Training. **A) soleus SSN, B) soleus PCSA, C) soleus wet weight, D) MG SSN, E) MG PCSA and F) MG wet weight** following eccentric training in male rats (n=12). The untrained (right leg) values are shown as blue lines and trained (left leg) values are shaded blue. Statistical comparisons were performed using a one-way repeated measures ANOVA (Training). Data are displayed as mean ± standard deviation. * Denotes significant differences.

### Concentric Training Mechanical Results

Torque increased ∼18% at 90**°** and ∼12% at 70**°** with training (*p*<0.001; *η*^2^=0.704), with an effect of angle (*p*<0.001; *η*^2^=0.999) but no training × angle interaction (*p*=0.460; *η*^2^=0.028; Figure 7). For passive tension there was an effect of training (*p*=0.039; *η*^2^=0.196; Figure 7) owing to an ∼37% and ∼14% increase in passive tension at 90**°** and 70**°** respectively. There was an effect of angle (*p*<0.001; *η*^2^=0.786), but no interaction of training × angle (*p*=0.953; *η*^2^=0). There was a non-significant increase in torque at peak power (*p*=0.188; *η*^2^=0.167) and a significant decline in angular velocity at peak power (*p*=0.008; *η*^2^=0.521; Supplemental Figure 4). Subsequently power was unchanged (*p*=0.076; *η*^2^=0.281; Figure 7).

**Figure 7.**
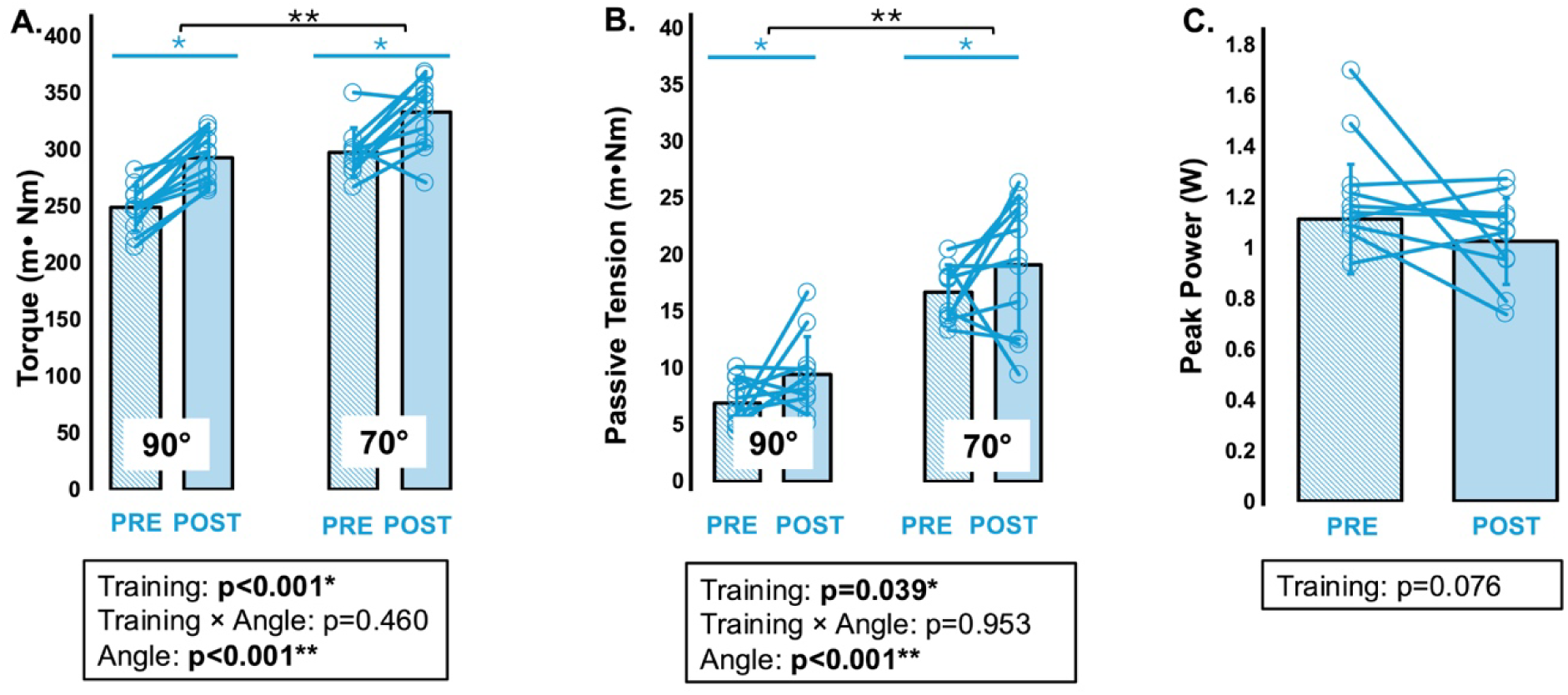
Concentric Training. **Plantar flexor A) Torque at 90° (shorter muscle length) and 70° (longer muscle length), B) 70 Passive tension at 90° and 70° and C) Peak power** following concentric training (n=11). Pretraining is shown as blue lines and post training is shown as shaded blue. Statistical comparisons were performed using a one-way repeated measures ANOVA (Training). Data are displayed as mean ± standard deviation. * Denotes significant differences.

### Concentric Training Morphological Results

There was no increase in soleus SSN following concentric training (∼2%; *p*=0.346; *η*^2^=0.081; Figure 8) with no change in FL or SL (Supplemental Figure 3). Additionally, there was no change in PCSA (*p*=0.918; *η*^2^=0.001; Figure 8) or wet weight (*p*=0.868; *η*^2^=0.003; Figure 8). MG SSN did not change with training (*p*=0.283; η^2^=0.104; Figure 8), nor did PCSA or wet weight, respectively (*p*=0.239; *η*^2^=0.124; Figure 8, *p*=0.819; *η*^2^=0.005; Figure 8).

**Figure 8.**
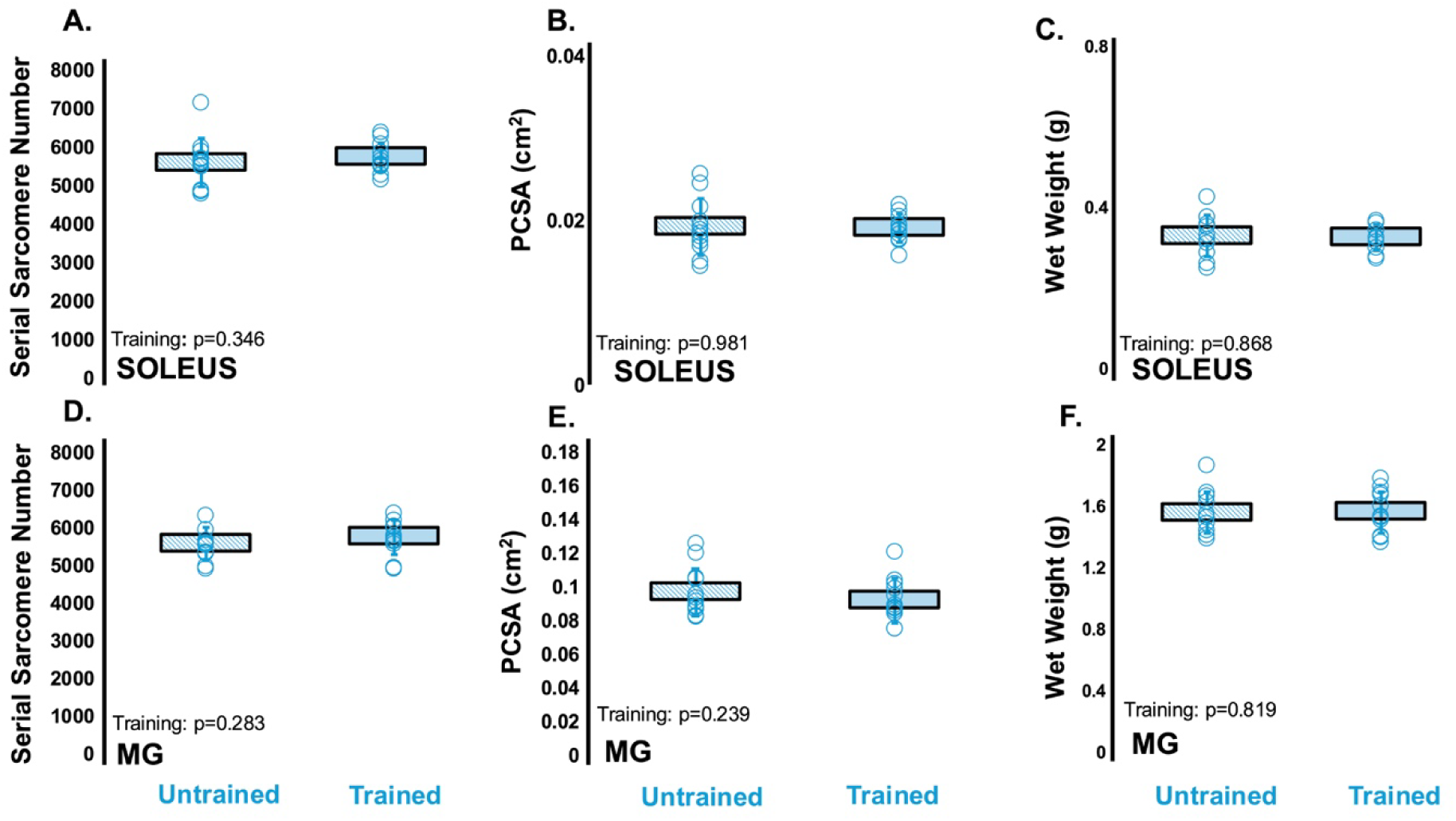
Concentric Training. **A) soleus SSN, B) soleus PCSA, C) soleus wet weight, D) MG SSN, E) MG PCSA and F) MG wet weight** following concentric (n=12) training. The untrained (right leg) values are shown as blue lines and trained (left leg) values are shaded blue. Statistical comparisons were performed using a one-way repeated measures ANOVA (Training). Data are displayed as mean ± standard deviation. * Denotes significant differences.

#### Training Modality Comparisons

##### Length dependent torque values achieved during training

As force at long muscle lengths is a potent loading parameter for SSN addition, to investigate torque at the longest muscle length (70° joint angle) achieved throughout the 3 training interventions, a representative rat with peak isometric torque closest to the mean across all training weeks was selected from each group. The average torque of set 1 across the 6 weeks was 218.3 ± 39.4, 188.0 ± 72.2 and 237.1 ± 31.9 mN·m for the isometric, eccentric, and concentric training, respectively. Meanwhile, the average peak torque for the first contraction of set #1 was 273.0 ± 43.3, 282.7 ± 108.6 and 283.9 ± 34.4 mN·m for isometric, eccentric. and concentric training. Therefore, peak torque at the longest MTU lengths during training was in a similar range across the 3 training modes.

##### Mechanical Adaptations

After adjusting for baseline differences, torque increased across all training groups (*p*=0.904; *η*^2^=0.003; Figure 9) and demonstrated an effect of angle (*p*=0.014; *η*^2^=0.092) owing to greater torque at longer muscle lengths (70°). There was no group × angle interaction (*p*=0.605; *η*^2^=0.016). Peak power did not differ between training groups (p=0.203; *η*^2^=0.098; Figure 10).

**Figure 9.**
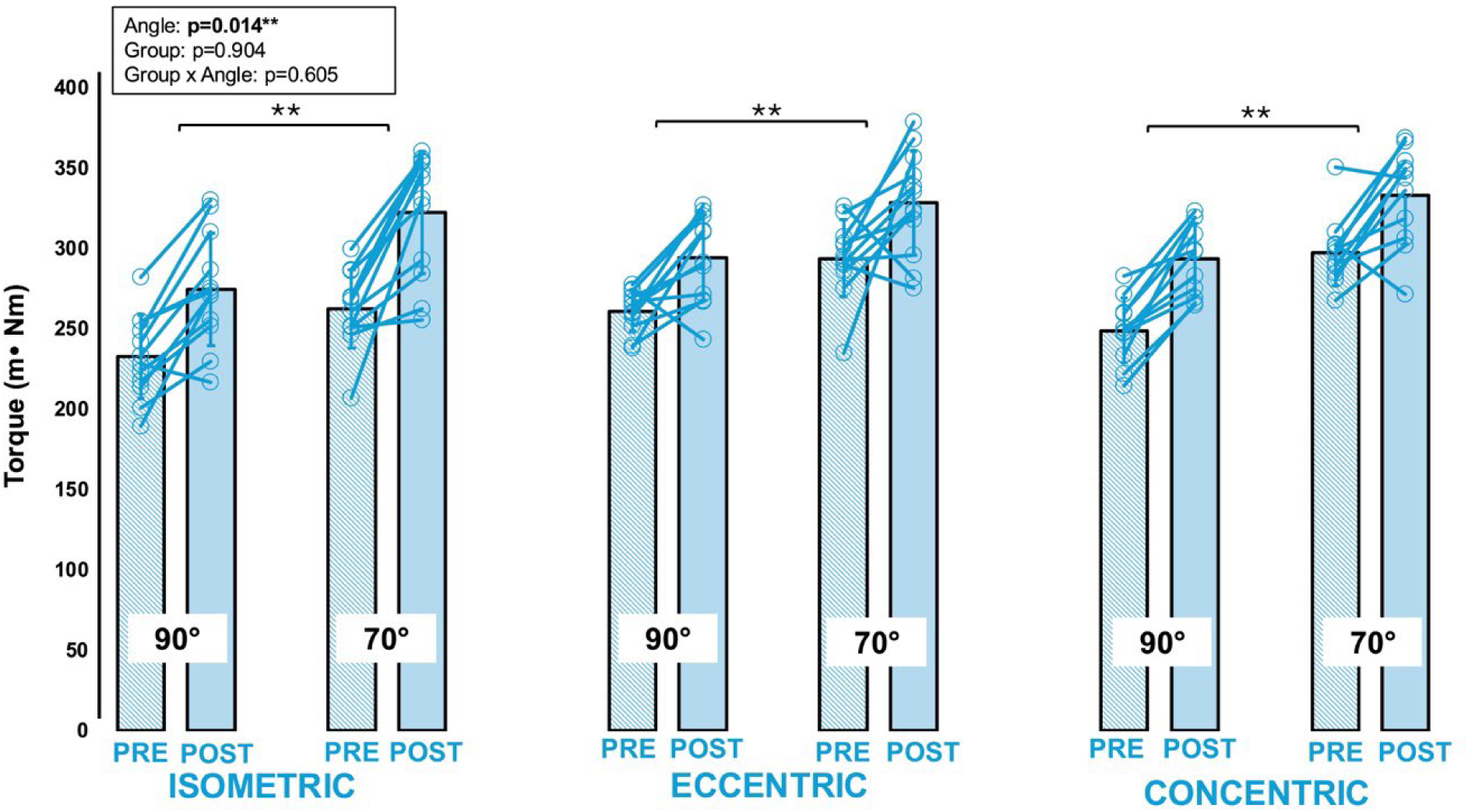
Increases in **Active torque** in response to isometric (n=12), eccentric (n=12) or concentric (n=11) training. Blue lines represent pre-training, and shaded blue is post-training. Statistical analysis was performed using an ANCOVA *Denotes statistical significance.

**Figure 10.**
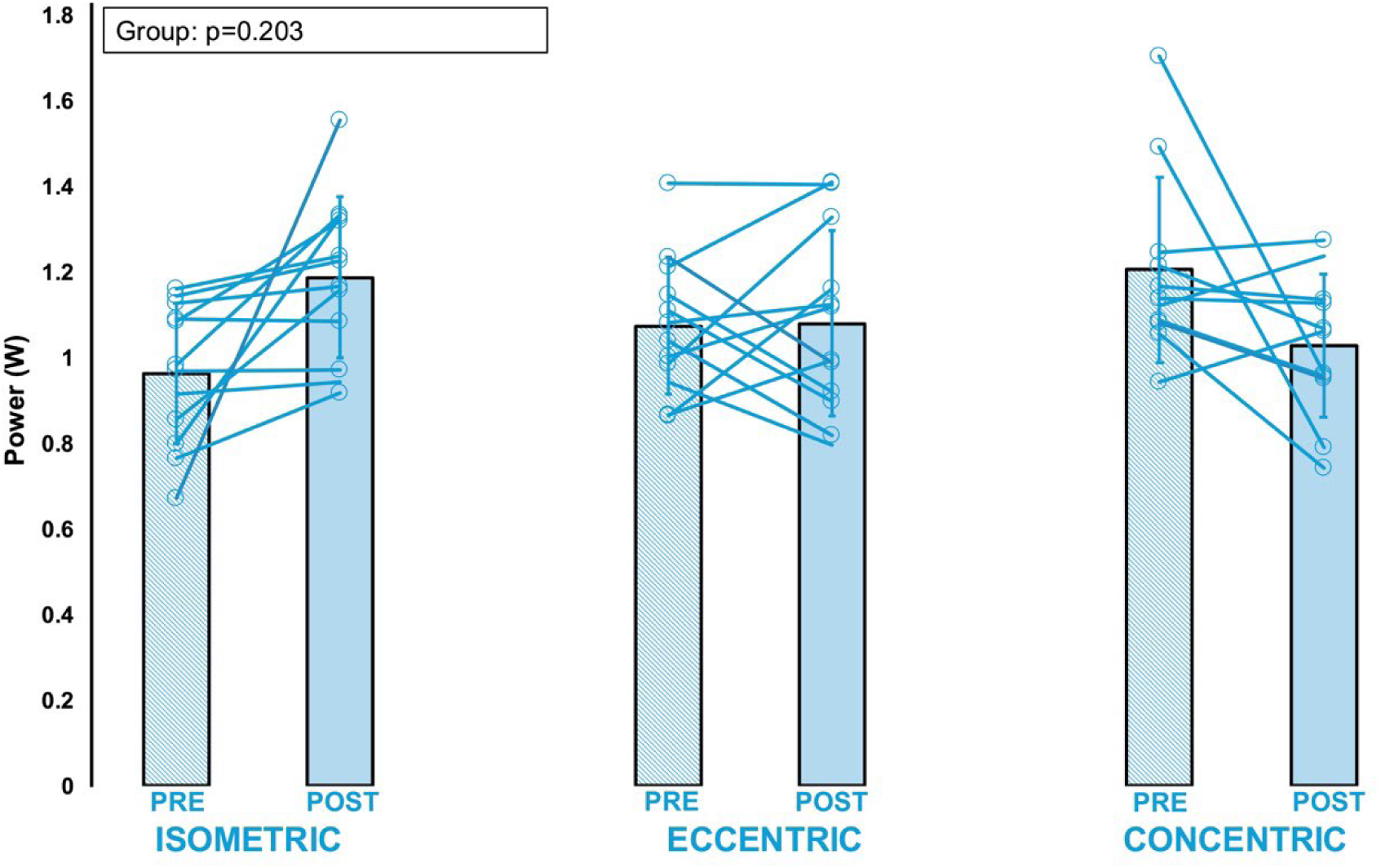
**Peak power (W)** with isometric (n=12), eccentric (n=12) or concentric (n=11) training in male rats. Blue lines represent pre-training, and shaded blue is post-training. Statistical analysis was performed using an ANCOVA. *Denotes statistical significance.

##### Morphological Adaptations-Soleus

After accounting for baseline differences in SSN, soleus SSN had an effect of training group (*p*=0.010; *η*^2^=0.249; Figure 11) where isometric (*p*=0.004) and eccentric (*p*=0.022) training increased SSN greater than concentric training, with no difference in adaptation between isometric and eccentric training (*p*=0.500). Soleus PCSA differed between training groups (*p*=0.044; *η*^2^*=*0.177; Figure 12) as concentric training yielded greater PCSA. Soleus wet weight did not differ between training groups (*p*=0.129; *η*^2^*=*0.120; Figure 13).

**Figure 11.**
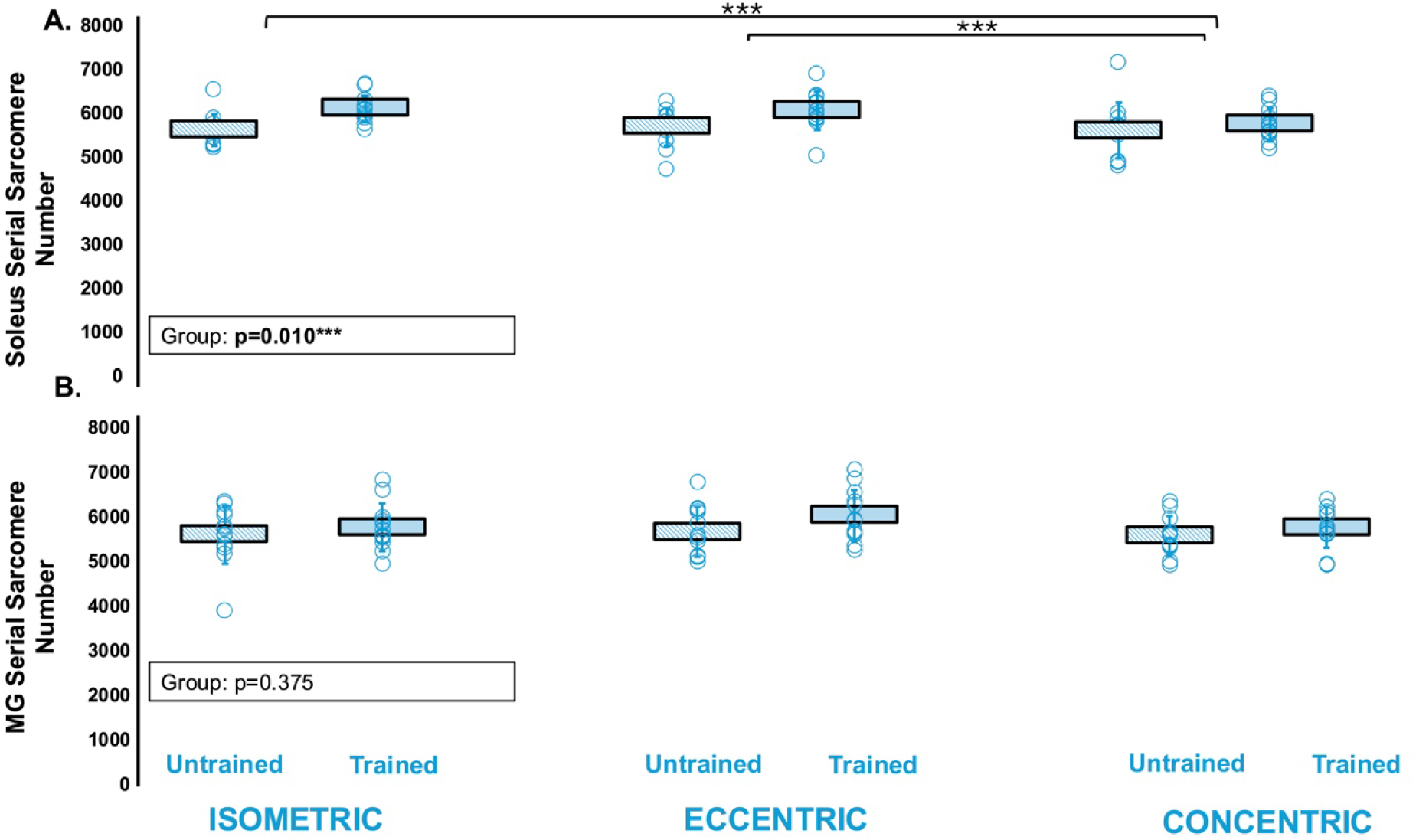
**A) soleus SSN and B) MG SSN** following isometric (n=12), eccentric (n=12) and concentric training (n=12). The untrained (right leg) is shown as blue lines and the trained (left leg) is shaded blue. Statistical analysis was performed using an ANCOVA. *Denotes significant differences.

**Figure 12.**
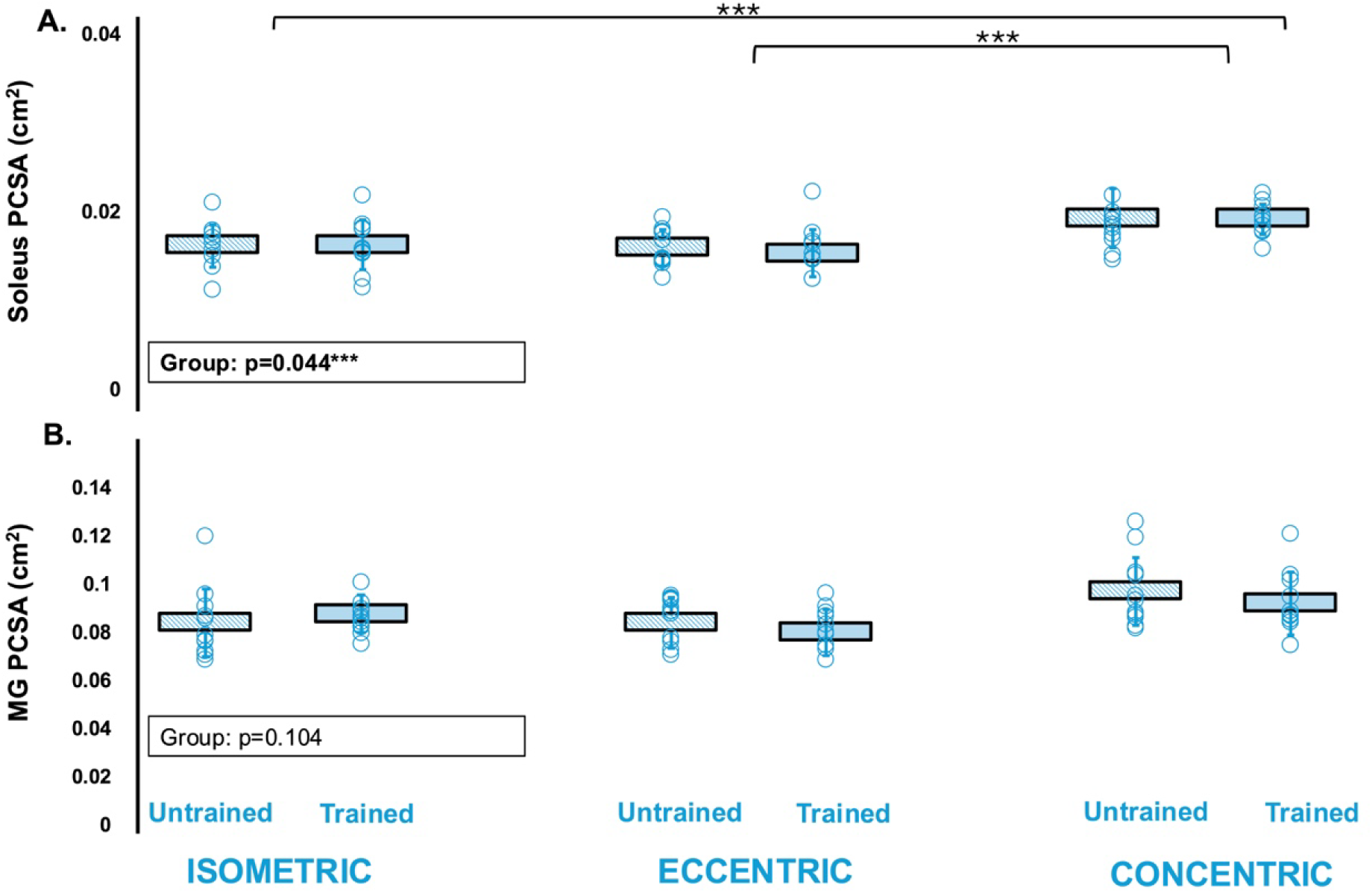
**A) soleus PCSA and B) MG PCSA** following isometric (n=12), eccentric (n=12) or concentric training (n=12). The untrained (right leg) is shown in shaded blue and the trained (left leg) is shown in blue. Statistical comparisons were performed using an ANCOVA. Data are displayed as mean ± standard deviation. * Denotes significant differences.

**Figure 13.**
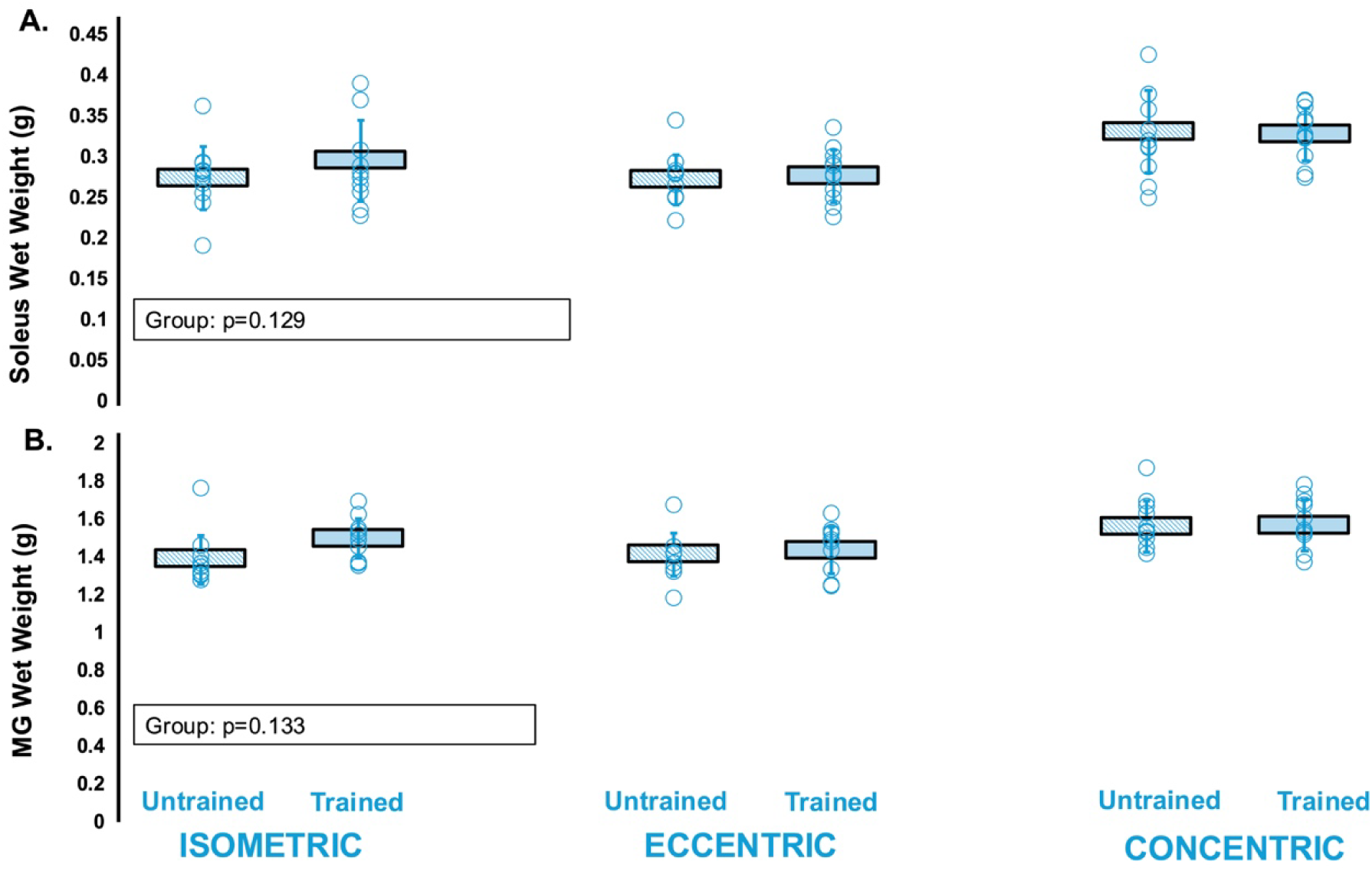
**A) soleus wet weight and B) MG wet weight** following isometric (n=12), eccentric (n=12) or concentric training (n=12). The untrained (right leg) is shown in shaded blue and the trained (left leg) is shown in blue. Statistical comparisons were performed using an ANCOVA. Data are displayed as mean ± standard deviation.

##### Morphological Adaptations-MG

After accounting for baseline differences in SSN, there was no effect of training group (*p*=0.375 *η*^2^=0.060; Figure 11) for MG SSN. MG PCSA and wet weight were not different between training groups (*p*=0.104; *η*^2^*=*0.132; Figure 12, *p=*0.133*; η*^2^*=*0.119; Figure 13, respectively).

## Discussion

In this study, we characterized contraction-type dependent remodelling of longitudinal muscle growth, specifically serial sarcomere addition following 4 and 6 weeks of resistance training. SSN adaptations are rapid and prioritized over radial growth therefore we collapsed the 4 and 6 week timepoints for analysis^18^. Using a dynamometer to standardize joint excursion, activation timing, and velocity, we found that both maximal isometric and submaximal eccentric training (60-70% of maximal isometric torque) produced similar increases in SSN (+7-9%). Notably, maximal concentric contractions failed to increase SSN despite being initiated at the same muscle lengths used during isometric training and at the muscle lengths reached at the end of eccentric contractions. Here we highlight that the regulation of longitudinal muscle remodelling is contraction-type dependent likely owing to the mechanical loading environments specific to eccentric training and isometric training at a longer muscle length.

### Isometric and eccentric training increased SSN in the soleus muscle

Based on some recent discussions in the literature^4–6^, we set out to address 3 testable hypotheses. Contrary to hypothesis i) *we should expect the largest magnitude of SSN increases following isometric training at longer MTU lengths,* and ii) *we should not expect an addition of SSN following the eccentric training intervention*, we observed the largest magnitude of SSN increases for both isometric training (+9%) and eccentric training (+7%) with no differences between training modalities (Figure 11), and for the MG the only increase in SSN was observed following eccentric training. Therefore, our findings indicate that not only end-contraction fibre length and high forces drive serial sarcomerogenic adaptations following resistance training, but likely additional stimuli throughout the active lengthening phase of the eccentric contraction also regulate the extent of serial sarcomerogenesis.

Isometric and eccentric training both involved muscle activation at the plateau region and nearing the descending limb of the force-length relationship (∼2.6 µm)^19^ corresponding to an intermediate muscle length. Isometric training was performed maximally, whereas, during eccentric contractions the muscle was lengthened to this same intermediate length, albeit during submaximal activation (60-70% of the current used to evoke maximal isometric torque). Given both contraction types resulted in similar torque production at the same final muscle length during training, then one could assume that fascicle/fibre/sarcomere lengths at the end of the eccentric contraction were likely similar to those of isometric. Notably, both isometric and eccentric training modalities arrived at a similar magnitude increase in SSN (Figure 11). The magnitude of sarcomerogenesis following both training modalities, showed a similar increase as observed by Ucar *et al*. (2023)^7^ during isometric training to intermediate muscle lengths (+∼12%), and our eccentric training adaptations were similar to Hinks *et al.* (2024) (+∼11%)^12^ with submaximal eccentric training.

In addition to active muscle lengthening, the mechanical loading environment that eccentric training creates highlights its effectiveness in promoting longitudinal muscle growth in humans^20,21^ and animal models^4,10,12,15^. During states of activity (i.e., not casted/immobilized) while high force/strain at long MTU lengths may be one of the primary stimuli of sarcomerogenesis, eccentric contractions offer a context-dependent stimulus for increases in SSN. Specifically, eccentric contractions combine high force production at long muscle lengths with substantial relaxion strain during de-activation^22^. For a MTU with a compliant tendon, throughout a lengthening muscle contraction (eccentric contraction) muscle-tendon unit compliance can allow fascicles to shorten against a compliant tendon as force increases. At the end of the eccentric phase and during the isometric hold, fascicle length stabilizes and then increases upon relaxation^23^. Notably, the magnitude of this post-contraction lengthening or relaxation strain has been strongly associated with training-induced increases in SSN and is a proposed potent stimulus for serial sarcomerogenesis^24^. It is also important to note that rodent tendons are relatively stiff compared with humans, so this effect of relaxation strain on SSN may indeed differ across species and amount of force generated. Thus, eccentric exercise indeed promotes increases in SSN as it drives the mechanotransduction signals regulating sarcomerogenesis. This distinction reflects differences in the underlying mechanisms governing sarcomerogenesis rather than the stimulus *per se*.

Moreover, evidence from our work on eccentric overload training indicates that added stress and strain during active lengthening are indeed key stimuli for SSN addition. In our previous work, both conventional resistance training (eccentric and concentric contractions at a current which evokes 60% of maximal isometric torque) and eccentric overload training (eccentric contractions at a current which evokes 100% of maximal isometric torque and concentric contractions at a current which evokes 60% of maximal isometric torque) groups experienced identical isometric pre-activation (500ms) at longer muscle lengths, yet the group performing eccentric overload produced greater increases in soleus SSN. Therefore, the additional stress and strain imposed during active lengthening likely contributed to longitudinal remodelling. These findings indicate that while isometric contractions remain an effective training modality for promoting increases in SSN, there may be additional mechanotransduction signals driven by eccentric contractions that promote serial sarcomerogenesis.

The third testable hypothesis presented was also refuted. iii) *Provided concentric contractions are initiated with high forces at longer muscle lengths, SSN should increase following concentric training*. Our results demonstrated no change in SSN with concentric training despite pre-activation (500ms) at the plateau nearing descending limb of force-length relationship (same muscle length as isometric and ending eccentric contraction length). These findings indicate that isometric pre-activation alone was insufficient to stimulate serial sarcomerogenesis, or to compensate for the loss of sarcomeres during active shortening. However, our work is the first to examine dynamometer controlled concentric training. Previous work on concentric biased exercise (uphill treadmill running) has shown a non-significant ∼2% decrease in soleus SSN^8^, a 5% decrease^17^, and no change in SSN^16^, suggesting a high degree of variability dependent on the muscle group and the extent of excursion.

### The biarticular MG only increased SSN with eccentric training

The soleus muscle had greater increases in SSN compared to the MG. The differing response between the soleus and MG is likely owing to the biarticular nature of the MG such that it crosses both the ankle and the knee joint ^25^. In the present study, the knee angle was at approximately 150°, a non-straight position, thus limiting excursion and disadvantaging the mechanical contribution of the MG muscle to plantar flexion. That said, the MG had the most robust (∼7%) increase in SSN in the eccentric, trained group compared to no change in the isometric and concentric groups (Figure 6). The MGs increase in SSN following eccentric training suggests that fibre stretch during active lengthening, independent of the initial muscle length, may itself provide a unique stimulus for sarcomerogenesis that was not present during other contraction modes. Furthermore, due to the longitudinal orientation of soleus fibres, changes in muscle length are transmitted more directly along the fibre axis which could lead to greater longitudinal mechanical strain, and a stronger mechanotransducive stimulus for serial sacromerogenesis. Conversely, the greater pennation of fibre orientation in the MG lowers the magnitude of longitudinal fibre strain. Therefore, differences in architecture between muscles would influence the mechanical stimulus for serial sacromerogenesis and contribute to the observed differences in adaptive responses.

### Similar changes in mechanical adaptations with training

Torque increased similarly for all training modalities despite differences in serial sarcomerogenesis and wet weight adaptations. The plantar flexors are composed of the soleus, medial gastrocnemius, lateral gastrocnemius, and plantaris. Therefore, increases in radial hypertrophy of the lateral gastrocnemius and plantaris may have supported increased torque production. The increases in torque observed in the present study differ from those observed by Lac and Cavalie *et al.* (1999) with 5 weeks of maximal isometric training (+36% maximal tetanic contraction)^26^, although, are similar to our lab’s previously observed +∼13% following 4 weeks of maximal eccentric contractions^27^. However, peak power increased 23% with isometric training with no change following eccentric or concentric training. Isometric training trended toward increasing soleus wet weight and MG wet weight (Figure 4), suggesting increases in radial growth may have contributed to enhanced power production^28^. Additionally, passive tension increased similarly across groups. In theory, the addition of sarcomeres in series should relieve passive tension by causing optimal SL to occur at a more stretched MTU length^29^. However, the equivocal increase across groups suggests there was likely a trade-off in which passive tension relieved through SSN addition^29^ was counteracted by extracellular matrix (ECM) remodelling. ^30, 31^ Heinemeier *et al.* (2007), compared extracellular matrix remodelling in isometric, eccentric and concentric trained rats and found COL1A1 mRNA expression (encodes type 1 collagen in the ECM) was increased only in the isometric and eccentric trained rat’s gastrocnemius after training^32^.

### Methodological considerations

As this study was longitudinal, rat maturation is a consideration. Growth related changes could have been a confounding factor to longitudinal muscle measures. However, we reduced this effect by ensuring rats had reached sexual maturity and maturation was no longer rapid^33^, and all morphological comparisons are within the same rat. The untrained right leg acted an internal control to the trained left leg^15,27,34^. Both legs matured similarly allowing the differentiation of training-induced changes versus maturation.

Additionally, given the importance of fascicle length/strain in regulating SSN addition, future work would benefit from having direct measures of not only MTU lengths, but relative contributions of muscle and tendon elongation to MTU, fascicle dynamics and matching activation during training to better understand the mechanical loading environment imposed by each distinct training modality. Future work can also analyze tendon remodelling because of training. Additionally, due to mechanical constraints of our experimental set-up, our range of motion was limited, and we were unable to construct a complete torque-angle relationship. Therefore, our assessments at 90° and 70° plantar flexion likely only represent the plateau near the descending limb^19^. Our in vivo mechanical testing is influenced by several synergistic plantar flexors which is not exclusive to the soleus and medial gastrocnemius in which morphological measures have been performed. Synergistic muscles and MTU compliance affect the theoretical, and canonical relationship between SSN and mechanical function. Therefore, future experiments may consider additional testing of ex-vivo mechanical function of individual muscle groups to better represent these relationships.

A further consideration is that during hindlimb fixation in 10% phosphate-buffered formalin, the ankle angle was fixed at 90° rather than at optimal muscle length (L_0_) for force production. Measuring both FL and SL at a standardized joint position allows for SSN to be estimated as a ratio of the FL to SL, which accounts for differences in SL between groups. Although the fixed joint position may result in muscle or group specific differences in SL due to variation in resting architecture or L0, these differences are incorporated into the calculation and therefore do not inherently confound the estimate of SSN.

## Conclusion

Isometric and eccentric training resulted in greater increases in soleus longitudinal remodelling through the addition of sarcomeres in series compared to concentric training. These findings highlight that the mechanical loading environment can augment different serial sarcomerogenic responses. Knowledge of contraction type dependence on muscle remodeling is critical when making targeted interventions to mitigate age-related sarcomere loss^1^ or establishing training or rehabilitation programs for athletes.

## Methods

### Animals

Thirty-six (n=18; 4-week timepoint, n=18; 6-week timepoint; divided evenly across 3 training groups) male Sprague-Dawley rats (mass=∼468g) ranging in age from 14-week-old to 18-week-old were obtained (Charles River Laboratories, QC, Canada). Rats were housed in groups of three at 23°C and given a Teklad global 18% protein rodent diet (Envigo, Huntington, Cambs., UK) and water *ad-libitum.* All protocols were approved by the University of Guelph’s Animal Care Committee (AUP #4905). Baseline pre-training mechanical testing was completed at 14 weeks to 18 weeks of age, then training commenced ∼3 days later, occurring 3 days/week for a total of 12 sessions over 4 weeks or 18 sessions over 6 weeks. Rodents have a ∼7-fold greater metabolic rate than humans ^35^, making this training program comparable to 4-6 months in humans. Post-training mechanical testing was conducted 48-72 hours following the final training session. All training and mechanical testing (i.e., pre-post-values) was conducted on the left leg with the right leg acting as an internal control for muscle wet mass, physiological cross-sectional area (PCSA), and SSN in accordance with previous studies ^12,36,37^. The soleus and medial gastrocnemius (MG) muscles were fixed then dissected for SSN determinations on both legs as described below ^12^.

### Mechanical Testing

Rats were anesthetized using inhaled isoflurane administered until a surgical plane of anaesthesia was reached (typically within 3–5 min) and maintained throughout all procedures until completion. Rats were then positioned in a supine position on a heated platform maintained at 37 °C. The left leg of the rat was shaved, and the leg was placed and secured to a force transducer/length controller foot pedal and fixed with tape (Figure 1b). Mechanical testing sessions began with determination of the stimulation current for maximal active plantar flexion torque (frequency = 100 Hz, pulse duration = 0.1 ms, train duration = 1 s) with a twitch before and after the tetanus contraction at an ankle angle of 90° (full plantar flexion = 180°). A 701C High-Powered, Bi-Phase Stimulator (Aurora Scientific, Aurora, ON, Canada) was used to evoke transcutaneous muscle stimulation, transmitted through the custom-made electrode ^38^ and used throughout the remainder of the testing session. A 100 Hz stimulation and a twitch pre/post tetanus was then completed at an ankle angle of 70°. Torque, angle, and stimulus trigger data were sampled at 1000 Hz with a 605A Dynamic Muscle Data Acquisition and Analysis System (Aurora Scientific, Aurora, ON, Canada) with a live torque tracing during all training and mechanical data collection sessions. To estimate the active torque, the baseline (passive torque) was subtracted from the maximum total torque value during stimulation ^39^. Passive torque at an ankle angle of 90° (shorter/neutral muscle length) and 70° (longer muscle length) was recorded to gain insight on adaptations in muscle passive properties. Passive torque was defined as the torque produced when the foot was clamped into the dynamometers at an ankle angle of 70° or 90°. Passive torque was assessed prior to active contractions. A torque-angular velocity-power relationship was established from isotonic contractions. Each isotonic contraction began at 70° and rotated to a joint angle of 110° against a fixed load clamp. The load clamps equated to 10%, 20%, 30%, 40%, 50%, 60%, 70%, and 80% of the maximum torque produced at 70°. The order of the isotonic contractions was randomized using an automated randomizer application. Angular velocity was recorded as the greatest slope of the position-time trace during shortening. Two minutes of rest separated each stimulation to minimize the development of muscle fatigue.

The estimated maximum shortening velocity at zero load (V_max_), maximum power (torque multiplied by angular velocity), and torque and velocity at peak power were determined by fitting the measured torque and angular velocity values to a rectangular hyperbolic curve ^39^.

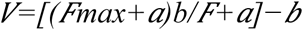

With F_max_ being maximum isometric torque at 70°, *F* and *V* representing the isotonic load clamp’s torque and angular velocity, respectively, and *a* and *b* representing Hill’s thermodynamic constants with the units of torque (mN·m) and velocity (°/s), respectively. All curve-fitting was performed in Python 3 using least square error optimization.

### Training programming

At the start of each session, the stimulus current was first adjusted to produce maximum isometric torque (pulse duration = 0.1ms, frequency = 100Hz, train duration = 500ms) at an ankle angle of 90° which ranged from 50 to 90 mA for all training modalities.

#### Isometric Resistance Training

The current used to evoke maximal isometric torque was used throughout the isometric training session. The training consisted of 3 (weeks 1-4) to 4 (weeks 5-6) sets of 8-11 repetitions where each repetition had: 1) a 2000-ms isometric activation at 70° (MG sarcomere length ∼2.6 µm)^19^, and 2) 2 seconds of rest before the next repetition. Two minutes of rest between sets was provided.

#### Isokinetic Concentric Resistance Training

The current to evoke maximal isometric torque was used throughout the concentric training session. The training consisted of 3 sets of 8-10 repetitions where each repetition had: 1) a 500-ms isometric pre-activation at 70°, 2) shortening from 70° (MG sarcomere length ∼2.6 µm)^22^ to 110° (MG SL ∼2.1 µm)^19^ at 40 (weeks 1-3) and 80°/s (weeks 4-6), and 3) 2 second rest before the next repetition. Two minutes of rest between sets was provided.

#### Isokinetic Eccentric Resistance Training

The current to evoke maximal isometric torque was established then adjusted to produce an isometric torque that as closely as possible matched 60% of maximum isometric torque. This 60% submaximal current was used throughout the eccentric training session for weeks 1-2, then increased to 70% for the remainder of training. Previous work from our lab showed this eccentric training contraction intensity increased SSN ^9^. The training consisted of 3 sets of 8-11 repetitions where each repetition had: 1) a 500-ms isometric pre-activation at 110°, 2) lengthening from 110° (MG SL ∼2.1 µm)^22^ to 70° (MG sarcomere length ∼2.6 µm)^19^ at 40 (week 1-3) and 80°/s (weeks 4-6), 3) 2 second rest before the next repetition. Two minutes of rest between sets was provided.

### Serial sarcomere number determination

Following their final mechanical testing session, rats were sacrificed via isoflurane anesthetization followed by CO_2_ asphyxiation and cervical dislocation. The hindlimbs were amputated, skinned and fixed in 10% phosphate-buffered formalin with the ankle and knee joints pinned at 90°. After fixation for 1-2 weeks, the soleus and MG were dissected off the lower leg, weighed, then re-submerged in formalin until the commencement of SSN estimations.

To commence the process of architectural measurements, the muscles were rinsed with phosphate-buffered saline, then digested in 30% nitric acid for 6-8 h to remove connective tissue and allow for individual muscle fascicles to be teased out^13^.

For each muscle, two fascicles were obtained from each of the proximal, middle, and distal regions (i.e., n = 6 fascicles total per muscle). The SSN and fascicle length (FL) values were averaged across these six fascicles for the analysis and reporting of data. Dissected fascicles were placed on glass microslides (VWR International, USA), then FLs were measured using ImageJ software (version 1.53f, National Institutes of Health, USA) from pictures captured by a level, tripod-mounted digital camera, with measurements calibrated to a ruler in plane with the fascicles. Sarcomere length (SL) measurements were taken at n = 6 different locations proximal to distal along each fascicle via laser diffraction (Coherent, Santa Clara, CA, USA) with a 5-mW diode laser (∼1 mm beam diameter, 635 nm wavelength) and custom LabVIEW program (Version 2011, National Instruments, Austin, TX, USA), for a total of n = 36 SL measurements per muscle. For each fascicle, the six SL measurements were averaged to obtain a value of average SL. Given the laser diameter of ∼1 mm, one SL measurement itself represents an average of hundreds to thousands of SLs. Our total quantity of SL and FL measurements is consistent with previous studies ^10^. For each fascicle, the six SL measurements were averaged to obtain a single value of average SL.

Serial sarcomere number of each fascicle was calculated as:

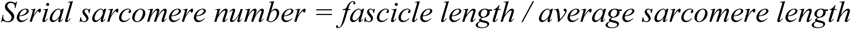

### Determination of physiological cross-sectional area

To gain further insight on changes in muscle contractile tissue in parallel, we calculated physiological cross-sectional area (PCSA; in cm^2^) using the equation ^40^:

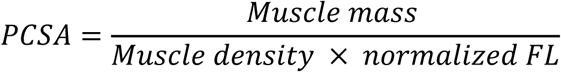

Muscle density was assumed to be 1.112 g/cm^3 41^. Normalized FL was calculated using FL of dissected fascicles and measured SL in the equation ^41^.

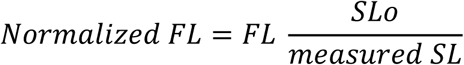

SLo represents optimal SL of rat muscle at rest, assumed to be ∼2.7 μm based on previous literature ^41^.

### Statistical analyses

All statistical analyses were performed in SPSS Statistics Premium version 28. Normality of data was confirmed using Shapiro-Wilk tests. There were no differences in SSN between 4 and 6 weeks, therefore data were collapsed across timepoint to increase statistical power. All statistical analyses listed below were carried out within each group; isometric, eccentric and concentric. Within group changes in SSN, SL, FL, PCSA and wet weight were assessed with a one-way analysis of variance (ANOVA) with repeated measures (Training [Trained leg, Untrained leg]). Changes in torque and passive tension were assessed with a two-way ANOVA with repeated measures (Training [Pre, Post] × Angle [90,70]. Differences in power, angular velocity at peak power and torque at peak power were assessed with a one-way ANOVA (Training [Pre, Post]. A Greenhouse-Geisser correction for sphericity was applied for all ANOVAs, and a Sidak correction was applied to all pairwise comparisons. Significance was set at α=0.05 and effect size is shown as (η^2^). To investigate differences between groups comparisons in power, SSN, SL, FL, muscle wet weight and PCSA between the trained and untrained leg were assessed by an Analysis of Covariance (ANCOVA) on post-values with Group as fixed factor and pre-values as the covariant. An ANCOVA was used to account for baseline differences between training groups. Differences in torque was assessed via an ANCOVA with fixed factors group and angle. A Greenhouse-Geisser correction for sphericity was applied for all ANCOVAs, and an LSD correction was applied to all pairwise comparisons. Significance was set at α=0.05 and effect size is shown as (η^2^).

## Acknowledgements

This project was supported by the Natural Sciences and Engineering Research Council of Canada (NSERC). We thank the Human Health Sciences Animal Care Staff for assistance with caring for our rats. We would like to thank Dr. Walter Herzog for comments on a previous version of this manuscript.

## Ethics statement

Approval was given by the University of Guelph’s Animal Care Committee and all protocols followed CCAC guidelines (AUP #4905).

## Data availability

All data generated or analysed during the study are available from the corresponding author upon request.

## Supplementary Information

Supplementary information is available for this paper.

## Author contributions

A.R., A.Z., A.H., and G.A.P. conceived and designed research; A.R., A.Z., K.C., A.K., M.S., performed experiments; A.R., A.Z., K.C., A.K., M.S., A.H., and G.A.P analyzed data; A.R., A.Z., K.C., A.K., M.S., A.H., and G.A.P interpreted results of experiments; A.R. prepared figures; A.R., A.Z., and G.A.P. drafted manuscript; A.R., A.Z., K.C., A.K., M.S., A.H., and G.A.P. edited and revised manuscript; A.R., A.Z., K.C., A.K., M.S., A.H., and G.A.P., approved final version of manuscript.

